# Haplotype-aware genotyping from noisy long reads

**DOI:** 10.1101/293944

**Authors:** Jana Ebler, Marina Haukness, Trevor Pesout, Tobias Marschall, Benedict Paten

**Author notes:** These authors contributed equally to this work (Jana Ebler, Marina Haukness, Trevor Pesout). Joint last/corresponding authors. Correspondence.

## Abstract

**Motivation:** Current genotyping approaches for single nucleotide variations (SNVs) rely on short, relatively accurate reads from second generation sequencing devices. Presently, third generation sequencing platforms able to generate much longer reads are becoming more widespread. These platforms come with the significant drawback of higher sequencing error rates, which makes them ill-suited to current genotyping algorithms. However, the longer reads make more of the genome unambiguously mappable and typically provide linkage information between neighboring variants.

**Results:** In this paper we introduce a novel approach for haplotype-aware genotyping from noisy long reads. We do this by considering bipartitions of the sequencing reads, corresponding to the two haplotypes. We formalize the computational problem in terms of a Hidden Markov Model and compute posterior genotype probabilities using the forward-backward algorithm. Genotype predictions can then be made by picking the most likely genotype at each site. Our experiments indicate that longer reads allow significantly more of the genome to potentially be accurately genotyped. Further, we are able to use both Oxford Nanopore and Pacific Biosciences sequencing data to independently validate millions of variants previously identified by short-read technologies in the reference NA12878 sample, including hundreds of thousands of variants that were not previously included in the high-confidence reference set.

## 1 Introduction

Reference-based genetic variant identification comprises two related processes: genotyping and phasing. Genotyping is the process of determining which genetic variants are present in an individual’s genome. A genotype at a given site describes whether both chromosomal copies carry a variant allele, only one of them carries it’ or whether the variant allele is not present at all. Phasing refers to determining an individual’s haplotypes, which consist of variants that lie near each other on the same chromosome and are inherited together. To completely describe all of the genetic variation in an organism, both genotyping and phasing are needed. Together, the two processes are called *diplotyping*.

Many existing variant analysis pipelines are designed for short DNA sequencing reads (1, 2). Though short reads are very accurate at a per-base level, they can suffer from being difficult to unambiguously align to the genome, especially in repetitive or duplicated regions (3). The result is that millions of bases of the reference human genome are not currently reliably genotyped by short reads, primarily in multi-megabase gaps near the centromeres and short arms of chromosomes (4). While short reads are unable to uniquely map to these regions, long reads can potentially span into or even across them. This makes it so long reads are advantageous over short reads for tasks such as haplotyping, large structural variant detection, and *de novo* assembly (5–8). Here, we attempt to demonstrate the utility of long reads for more comprehensive genotyping.

Long read DNA sequencing technologies are rapidly falling in price and increasing in general availability. Such technologies include Single Molecule Real Time (SMRT) Sequencing by Pacific Biosciences (PacBio) and nanopore sequencing by Oxford Nanopore Technologies (ONT). However, due to the historically greater relative cost and higher sequencing error rates of these technologies, little attention has been given thus far to the problem of genotyping single nucleotide variants (SNVs) with long reads. Recently, (9) have taken first steps in this direction, but their approach does not scale to process whole human genomes in reasonable time.

For an illustration of the benefit of using long reads to diplo-type, consider Figure 1. Shown are three SNV positions covered by long reads. The gray sequences represent the true haplotype sequences and reads are colored in blue and red. The colors correspond to the haplotype which the respective read stems from: the red ones from the upper sequence, and the blue ones from the lower one. Since sequencing errors can occur, the alleles supported by the reads are not always equal to the true ones in the haplotypes shown in gray. Considering the SNVs individually, we would probably genotype the first one as A/C, the second one as T/G and the third one as G/C, since the number of reads supporting each allele are the same. This leads to a wrong genotype prediction for the second SNV. However, if we knew which haplotype each read stems from, that is, if we knew their colors, then we would be unsure about the genotype of the second SNV. It could also be G/G or T/T, since the reads stemming from the same haplotypes must support the same alleles. Therefore, using haplotype information during genotyping makes it possible to compute more reliable genotype predictions and to detect uncertainties.

**Fig. 1.**
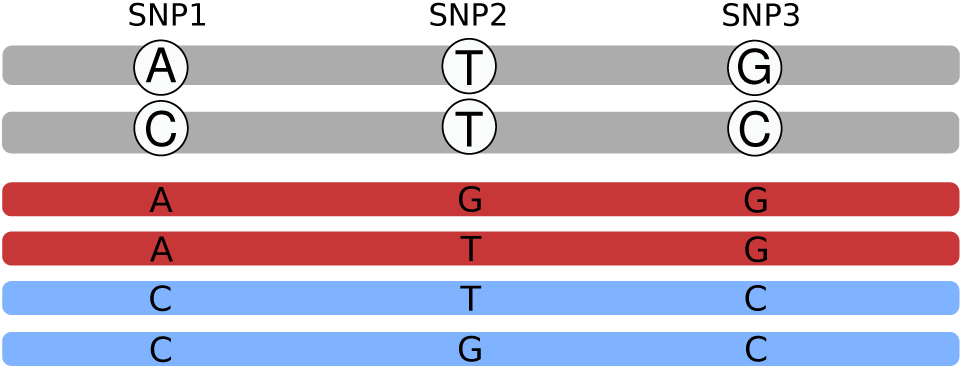
Motivation. Gray sequences illustrate the haplotypes; the reads are shown in red and blue. The red reads originate from the upper haplotype, the blue ones from the lower Genotyping each SNV individually would lead to the conclusion that all of them are heterozygous. Using the haplotype context reveals uncertainty about the genotype of the second SNV.

### Contributions

In this paper, we show that for contemporary long read technologies, read-based phase inference can be simultaneously combined with the genotyping process for SNVs to produce accurate \ and to detect variants in regions not mappable by short reads. We show that key to this inference is the detection of linkage relationships between heterozygous sites within the reads. To do this, we describe a novel algorithm to accurately predict diplotypes from noisy long reads that scales to deeply sequenced human genomes. We achieve this by considering bipartitions of all given sequencing reads, corresponding to the two haplotypes of an individual. The problem is formalized using a Hidden Markov Model (HMM) from which we compute genotype likelihoods using the forward-backward algorithm and make genotype predictions by determining the likeliest genotype at each position.

We then apply this algorithm to diplotype one individual from the 1000 Genomes Project, NA12878, using long reads from both PacBio and ONT. NA12878 has been extensively sequenced and studied, and the Genome in a Bottle consortium has published sets of highly confident variant calls (10). We demonstrate that our method is accurate, that it can be used to confirm variants in regions of uncertainty, and that it allows for the discovery of variants in regions which are unmappable using short DNA read sequencing technologies.

## 2 Methods

We describe a probabilistic model for diplotype and genotype inference, and in this paper use it to find maximum posterior probability genotypes. The approach builds upon the WhatsHap approach (10), but incorporates a full probabilistic allele inference model into the problem. It has similarities to that proposed by Kuleshov et al. (11), but we here frame the problem using Hidden Markov Models (HMMs).

### 2.1 Alignment Matrix

Let **M** be an alignment matrix whose rows represent sequencing *reads* and whose columns represent genetic *sites*. Let *m* be the number of rows, let *n* be the number of columns, and let **M**_*i,j*_ be the *j*th element in the *i*th row. In each column let ∑_*j*_ ⊂ ∑ represent the set of possible *alleles* such that **M**_*i,j*_ ∈ ∑_*j*_ ⋃ {–}, the “–” gap symbol representing a site at which the read provides no information. We assume no row or column is composed only of gap symbols, an uninteresting edge case. An example alignment matrix is shown in Figure 2. Throughout the following we will be informal and refer to a row *i* or column *j*, being clear from the context whether we are referring to the row or column itself or the coordinate.

**Fig. 2.**
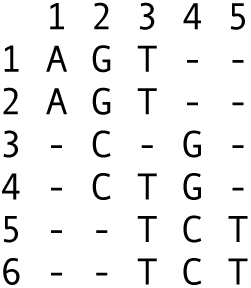
Alignment Matrix. Here, the alphabet of possible alleles is the set of DNA nucleotides, i.e. ∑ = {A, C, G, T}

### 2.2 Genotype Inference Problem Overview

A diplotype *H* = (*H*^1^,*H*^2^) is a pair of haplotype (segments); a *haplotype* (segment) 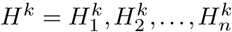 is a sequence of length *n* whose elements represents alleles such that 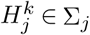. Let *B* = (*B*^1^, *B*^2^) be a bipartition of the rows of **M** into two parts (sets): *B*^1^, the first part, and *B*^2^, the second part. We use bipartitions to represent which haplotypes, of the two in a genome, the reads came from. By convention we assume that the first part of *B* are the reads arising from *H*^1^ and the second part of *B* are the reads arising from *H*^2^.

The problem we analyze is based upon a probabilistic model that essentially represents the (Weighted) Minimum Error Correction (MEC) problem (12, 13), while modeling the evolutionary relationship between the two haplotypes and so imposing a cost on bipartitions that create differences between the inferred haplotypes.

For a bipartition *B*, and making an i.i.d. assumption between sites in the reads:

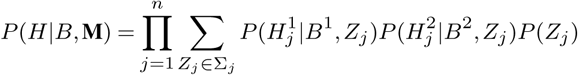

Here *P*(*Z*_*j*_) is the prior probability of the ancestral allele *Z*_*j*_ of the two haplotypes at column *j*, by default we can use a simple flat distribution over ancestral alleles (but see below). The posterior probability 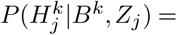

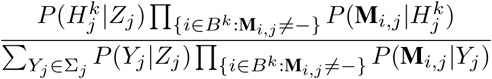

for *k* ∈ {1,2}, where the probability 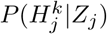 is the probability of the haplotype allele 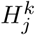 given the ancestral allele *Z*_*j*_. For this we can use a continuous time Markov model for allele substitutions, such as Jukes-Cantor (15), or some more sophisticated model that factors the similarities between alleles (see below). Similarly, 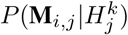 is the probability of observing allele **M**_*i,j*_ in a read given the haplotype allele 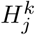.

The genotype inference problem we consider is finding for each site:

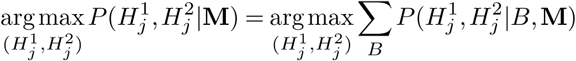

i.e. finding the genotype 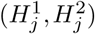 with maximum posterior probability for a generative model of the reads embedded in **M**.

### 2.3 A Graphical Representation Of Read Partitions

For a column *j* in M, a row *i* is *active* if the first non-gap symbol in row *i* occurs at or before column *j* and the last non-gap symbol in row *i* occurs at or after column *j*. Let 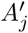 be the set of active rows of column *j*. For a column *j* a row *i* is *terminal* if its last non-gap symbol occurs at column *j* or *j* = *n*. Let *A’*_*j*_ be the set of active, non-terminal rows of column *j*.

Let 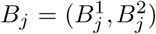 be a bipartition of *A*_*j*_ into a first part 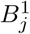 and a second part 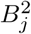. Let **B**_j_ be the set of all possible such bipartitions of the active rows of *j*. Similarly, let 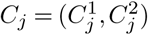 be a bipartition of 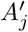, and *C*_*j*_ be the set of all possible such bipartitions of the active, non-terminal rows of *j*.

For two bipartitions *B* = (*B*^1^,*B*^2^) and *C* = (*C*^1^,*C*^2^),*B* is *compatible* with *C* if the subset of *B*^1^ in *C*^1^ ∪ *C*^2^ is a subset of *C*^1^, and, similarly, the subset of *B*^2^ in *C*^1^ ∪ *C*^2^ is a subset of *C*^2^. Note this definition is symmetric and reflexive, although not transitive.

Let *G* = (*V*_*G*_,*Eg*) be a directed graph. The vertices *V*_*G*_ are the set of bipartitions of both the active rows and the active, non-terminal rows for all columns of M and a special *start* and *end* vertex, i.e. *V*_*G*_ = {*start, end*} ∪ (∪_*j*_ **B**_j_ ∪ *C*_*j*_). The edges *E*_*G*_ are a subset of compatibility relationships, such that (1) for all *j* there is an edge (*B*_*j*_ ∈ **B**_**j**_,*C*_*i*_ ∈ **C**_**j**_) if *B*_*j*_ is compatible with *C*_*j*_,(2) for all 0 < *j* < *n* there is an edge (*C*_*j*_ ∈ *C*_*j*_,*B*_*j*+1_ ∈ **B**_j+1_) if *C*_*j*_ is compatible with *B*_*j*+1_, (3) there is an edge from the start vertex to each member of **B**_1_, and (4) there is an edge from each member of **B**_n_ to the end vertex (Note that **C**_n_ is empty and so contributes no vertices to *G*). Figure 3 shows an example graph.

**Fig. 3.**
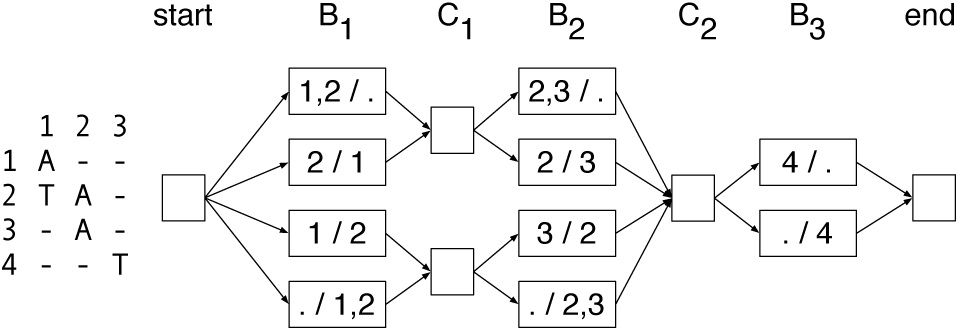
Example Graph. Left: An alignment matrix. Right: The corresponding directed graph representing the bipartitions of active rows and active non-terminal rows, where the labels of the nodes indicate the partitions, e.g. ‘1,2/.’ is shorthand for A =({1,2}, {}}).

The graph *G* has a large degree of symmetry and the following properties are easily verified:

- For all *j* and all *B*_*j*_ ∈ **B**_j_, the indegree and outdegree of *B*_*j*_ is 1.
- For all *j* the indegree of all members of **C**_**j**_ is equal.
- Similarly, for all *j* the outdegree of all members of **C**_**j**_ is equal.

Let the *maximum coverage*, denoted *maxCov*, be the maximum cardinality of a set *A*_*j*_ over all _*j*_. By definition, *maxCov* ≤ *m*. Using the above properties it is easily verified that: (1) the cardinality of *G* (number of vertices) is bounded by this maximum coverage, being less than or equal to 2 + (2*n* — 1)2^*maxCov*^, and (2) the size of *G* (number of edges) is at most 2*n*2^*maxCov*^.

Let a directed path from the start vertex to the end vertex be called a *diploid path*, *D* = (*D*_1_ = *start*,*D*_2_,…,*D*_2*n* +1_ = *end*). The graph is naturally organized by the columns of M, so that 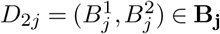 and 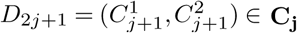 **C**_j_ for all 0 < *j* ≤ *n*. Let 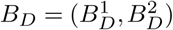 denote a pair of sets, where 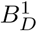 is the union of the first parts of the vertices of *D*_2_,…, *D*_2*n+*1_ and, similarly, 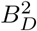 is the union of second parts of the vertices of *D*_2_,…,*D*_2*n+*1_.

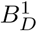 and 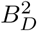 are disjoint because otherwise there must exist a pair of vertices within *D* that are incompatible, which is easily verified to be impossible. Further, because *D* visits a vertex for every column of M, it follows that the sum of the cardinalities of these two sets is *m*. *BD* is therefore a bipartition of the rows of M which we call a *diploid path bipartition*.

#### Lemma 1

The set of diploid path bipartitions is the set of bipartitions of the rows of M and each diploid path defines a unique diploid path bipartition.

*Proof:* We first prove that each diploid path defines a unique bipartition of the rows of M. For each column *j* of M, each vertex *B*_*j*_ ∈ **B**_**j**_ is a different bipartition of the same set of active rows. *B*_*j*_ is by definition compatible with a diploid path bipartition of a diploid path that contains it, and incompatible with every other member of **B**_**j**_. It follows that for each column *j* two diploid paths with the same diploid path bipartition must visit the same node in **B**_**j**_, and, by identical logic, the same node in **C**_**j**_, but then two such diploid paths are therefore equal.

There are 2^*m*^ partitions of the rows of M. It remains to prove that there are 2^*m*^ diploid paths. By the structure of the graph, the set of diploid paths can be enumerated backwards by traversing right-to-left from the end vertex by depth-first search and exploring each incoming edge for all encountered nodes. As stated previously, the only vertices with indegree greater than one are for all *j* the members of **C**_j_, and each member of **C**_j_ has the same indegree. For all *j* the indegree of *C*_*j*_ is clearly 2^|*C*_*j*_|-|*B*_*j*_|^: two to the power of the number of number of active, terminal rows at column *j*. The number of possible paths must therefore be 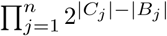. As each row is active and terminal in exactly one column, we obtain *m* = ∑_*j*_ |*C*_*j*_|−|*B*_*j*_| and therefore:

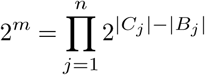

### 2.4 A Hidden Markov Model For Genotype and Diplotype Inference

In order to infer diplotypes, we define a Hidden Markov Model which is based on G, but additionally represents all possible genotypes at each genomic site (i.e. in each B column). To this end, we define the set of states **B**_**j**_ × ∑_*j*_ × ∑_*j*_, which contains a state for each bipartition of the active rows at position *j* and all possible assignments of alleles in ∑_*j*_ to the two partitions. Additionally, the HMM contains a hidden state for each bipartition in **C**_**j**_, exactly as defined for *G* above. Transitions between states are defined by the compatibility relationships of the corresponding bipartitions as before. This HMM construction is illustrated in Figure 4.

For all *j* and all *C*_*j*_ ∈ **C**_**j**_ each outgoing edge has transition probability *P*(*a*_1_***,a_2_***) = ∑(*a*_1_|*z*_*j*_)*P*(*a*_2_|*Z*_*j*_)*P*(*Z*_*j*_), where (*B*_*j*_,*a*_1_,*a*_2_) ∈ **B**_**j**_ × ∑_*j*_ × ∑_*j*_ is the state being transitioned to. Similarly, each outgoing edge of the start node has transition probability *P*(*a*_1_,*a*_2_). The outdegree of all remaining nodes is ***1***, so these edges have transition probability ***1***.

The start node, the end node, and members of **C**_**j**_ for all *j* are silent states, and hence do not emit symbols. For all *j*, members of **B**_**j**_ × ∑_*j*_ × ∑_*j*_ output the entries in the j-th column of **M** that are different from “-”. We assume every matrix entry to be associated with an error probability, which we can compute from 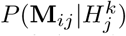 defined previously. Based on this, the probability of observing a specific output column of M can be easily calculated.

**Fig. 4.**
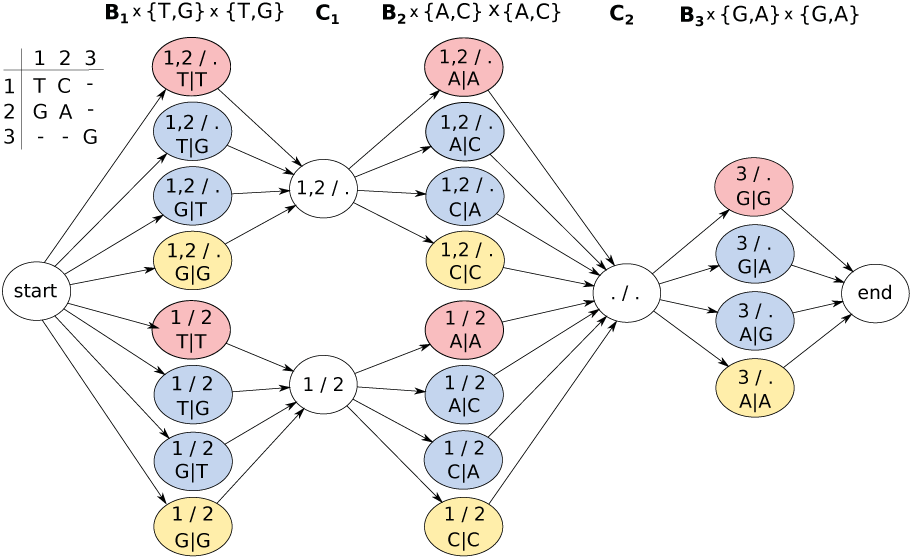
Genotyping HMM. Colored states correspond to bipartitions of reads and allele assignments at that position. States in C_1_ and C_2_ correspond to bipartitions of reads covering positions 1 and 2 or 2 and 3, respectively. In order to compute genotype likelihoods after running the forward-backward algorithm, states of the same color have to be summed up in each column.

#### 2.4.1 Computing Genotype Likelihoods

The goal is to compute genotype likelihoods for the possible genotypes for each variant position using the HMM defined above. Performing the forward-backward algorithm returns forward and backward probabilities of all hidden states. Using those, the posterior distribution of a state (*B*,*a*_1_,*a*_2_) ∈ **B**_**j**_ × ∑_*j*_ × ∑_*j*_, corresponding to bipartition B and assigned alleles *a*_1_ and *a*_2_, can be computed as

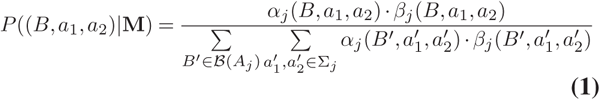

where *α*_*j*_(*B*,*a*_1_,*a*_2_) and *β*_*j*_(*B*,*a*_1_,*a*_2_) denote forward and backward probabilities of the state (*B,a*_1_*,a*_2_) and 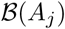, the set of all bipartitions of *A*_*j*_. The above term represents the probability for a bipartition *B* = (*B*^1^,*B*^2^) of the reads in *A*_*j*_ and alleles *a*_1_ and *a*_2_ assigned to these partitions. In order to finally compute the likelihood for a certain genotype, one can marginalize over all bipartitions of a column, and all allele assignments corresponding to that genotype.

##### Example 2.1

In order to compute genotype likelihoods for each column of the alignment matrix, posterior state probabilities corresponding to states of the same color in Figure 4 need to be summed up. For the first column, adding up the red probabilities gives the genotype likelihood of genotype *T/T,* blue of genotype *G/T* and yellow of *G/G*.

### 2.5 Implementations

We created two independent software implementations of this model, one based upon WhatsHap and one from scratch, which we call MarginPhase. Each uses different optimizations and heuristics that we briefly describe.

#### 2.5.1 WhatsHap Implementation

We extended the implementation of WhatsHap (11, bitbucket.org/whatshap/whatshap) to enable haplotype aware genotyping of bi-allelic variants based on the above model. WhatsHap focuses on re-genotyping variants, i.e. it assumes SNV positions to be given. In order to detect variants, a simple SNV calling pipeline was developed. It is based on samtools mpileup (16) which provides information about the bases supported by each read covering a genomic position. A set of SNV candidates is generated by selecting genomic positions at which the frequency of a non-reference allele is above a fixed threshold (0.25 for PacBio data, 0.4 for Nanopore data) and the absolute number of reads supporting the non-reference allele is at least 3.

##### Allele Detection

In order to construct the alignment matrix, a crucial step is to determine whether each read supports the reference or the alternative allele at each of *n* given genomic positions. In WhatsHap, this is done based on re-aligning sections of the reads (17). Given an existing read alignment from the provided BAM file, its sequence in a window around the variant is extracted. It is aligned to the corresponding region of the reference sequence and additionally, to the alternative sequence, which is artificially produced by inserting the alternative allele into the reference. The alignment cost is computed by using affine gap costs. Phred scores representing the probabilities for opening and extending a gap and for a mismatch in the alignment can be estimated from the given BAM file. The allele leading to a lower alignment cost is assumed to be supported by the read and is reported in the alignment matrix. If both alleles lead to the same cost, the corresponding matrix entry is “-”. The absolute difference of both a a phred scaled probability for the allele being wrong and is utilized for the computation of output probabilities.

##### Read Selection

Our algorithm enumerates all bipartitions of reads colignment scores is assigned as a weight to the corresponding entry in the alignment matrix. It can be interpreted asvering a variant position and thus has a runtime exponential in the maximum coverage of the data. To ensure that this quantity is bounded, the same read selection step implemented previously in the WhatsHap software is run before constructing the HMM and computing genotype likelihoods. Briefly, a heuristic approach described in (18) is applied, which selects phase informative reads iteratively taking into account the number of heterozygous variants covered by the read and its quality.

##### Transitions

Defining separate states for each allele assignment in **B**_**j**_ enables easy incorporation of prior genotype likelihoods by weighting transitions between states in **C**_j−1_ and **B**_j_ × ∑_*j*_ × ∑_*j*_. Since there are two states corresponding to a heterozygous genotype in the bi-allelic case (0|1 and 1|0), the prior probability for the heterozygous genotype is equally spread between these states.

In order to compute such genotype priors, the same likelihood function underlying the approaches described in (19) and (20) was utilized. For each SNV position, the model computes a likelihood for each SNV to be absent, heterozygous, or homozygous based on all reads that cover a particular site. Each read contributes a probability term to the likelihood function, which is computed based on whether it supports the reference or the alternative allele (19). Furthermore, the approach accounts for statistical uncertainties arising from read mapping and has a runtime linear in the number of variants to be genotyped (20). Prior genotype likelihoods are computed before read selection. In this way, information of all input reads covering a position can be incorporated.

#### 2.5.2 MarginPhase Implementation

MarginPhase (github.com/benedictpaten/marginPhase) is an experimental, open source implementation of the described HMM written in C. It differs from the WhatsHap implementation in the method it uses to explore bipartitions and the method to generate allele support probabilities from the reads.

##### Read Bipartitions

The described HMM scales exponentially in terms of increasing read coverage. For typical 20-60x sequencing coverage (i.e. average number of active rows per column) it is impractical to store all possible bipartitions of the rows of the matrix. MarginPhase implements a simple, greedy pruning and merging heuristic outlined in recursive pseudocode in Algorithm 1.

The procedure computePrunedHMM takes an alignment matrix and returns a connected subgraph of the HMM for M that can be used for inference, choosing to divide the input alignment matrix into two if the number of rows exceeds a threshold *t*, recursively.

The sub-procedure mergeHMMs takes two pruned HMMs for two disjoint alignment matrices with the same number of columns and joins them together in the natural way such that if at each site *i* there are 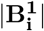 states in **HMM**_1_ and 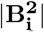 in **HMM**_2_ then the resulting HMM will have 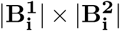 states. This is illustrated in Figure 5. In the experiments used here *t* = 8 and *v* = 0.01.

**Fig. 5.**
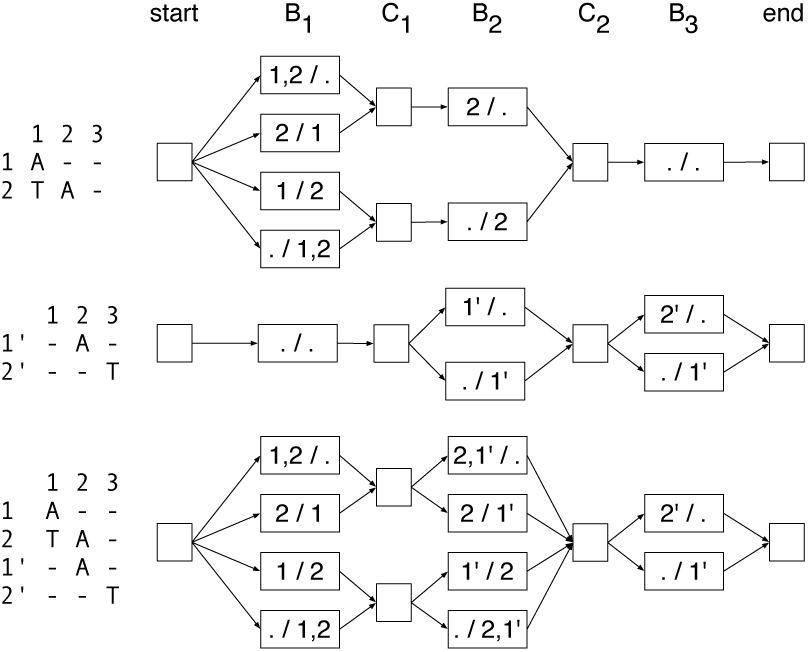
The merger of two read partitioning HMMs with the same number of columns. Top and middle: Two HMMs to be merged; bottom: the merged HMM. Transition and emission probabilities not shown.

##### Allele Supports

In MarginPhase the alignment matrix has a site for each base in the reference genome. To generate the allele support from the reads, for each read we calculate the posterior probability of each allele using the implementation of the banded forward-backward pairwise alignment described in (21). The result is that for each reference base, for each read that overlaps (according to an initial guide alignment extracted from the SAM/BAM file) the reference base we calculate the probability of each possible nucleotide (i.e. {‘A’, ‘C’, ‘G’, ‘T’}). Gaps are ignored and treated as missing data. This approach allows summation over all alignments within the band.

###### Algorithm 1

~~~
**procedure** COMPUTEPRUNEDHMM(M)
     **if** maxCov ≥ *t* **then**
                       Divide **M** in half to create two matrices, **M**_**1**_ and
                        **M**_**2**_, such that **M**_**1**_ is the first 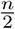 rows of **M**
                        and **M**_**2**_ is the remaining rows of **M**.
                 **HMM**_**1**_ ← computePrunedHMM(**M**_**1**_)
                 **HMM**_**2**_ ← computePrunedHMM(**M**_**2**_)
                 **HMM** ← mergeHMMs(**HMM**_**1**_, **HMM**_2_)
         else
             Let **HMM** be the read partitioning HMM for **M**.
   **return** subgraph of **HMM** including visited states
         and transitions each with posterior probability of
         being visited ≥ *v*, and which are on a path from
         the start to end nodes.
~~~

## 3 Results

### 3.1 Data Preparation and Evaluation

To test our methods, we used sequencing data for NA12878 from two different long read sequencing technologies. NA12878 is a participant from the 1000 Genomes Project (2) who has been extensively sequenced and analyzed. We used Oxford Nanopore reads from (7) and PacBio reads from (26).

**Fig. 6.**
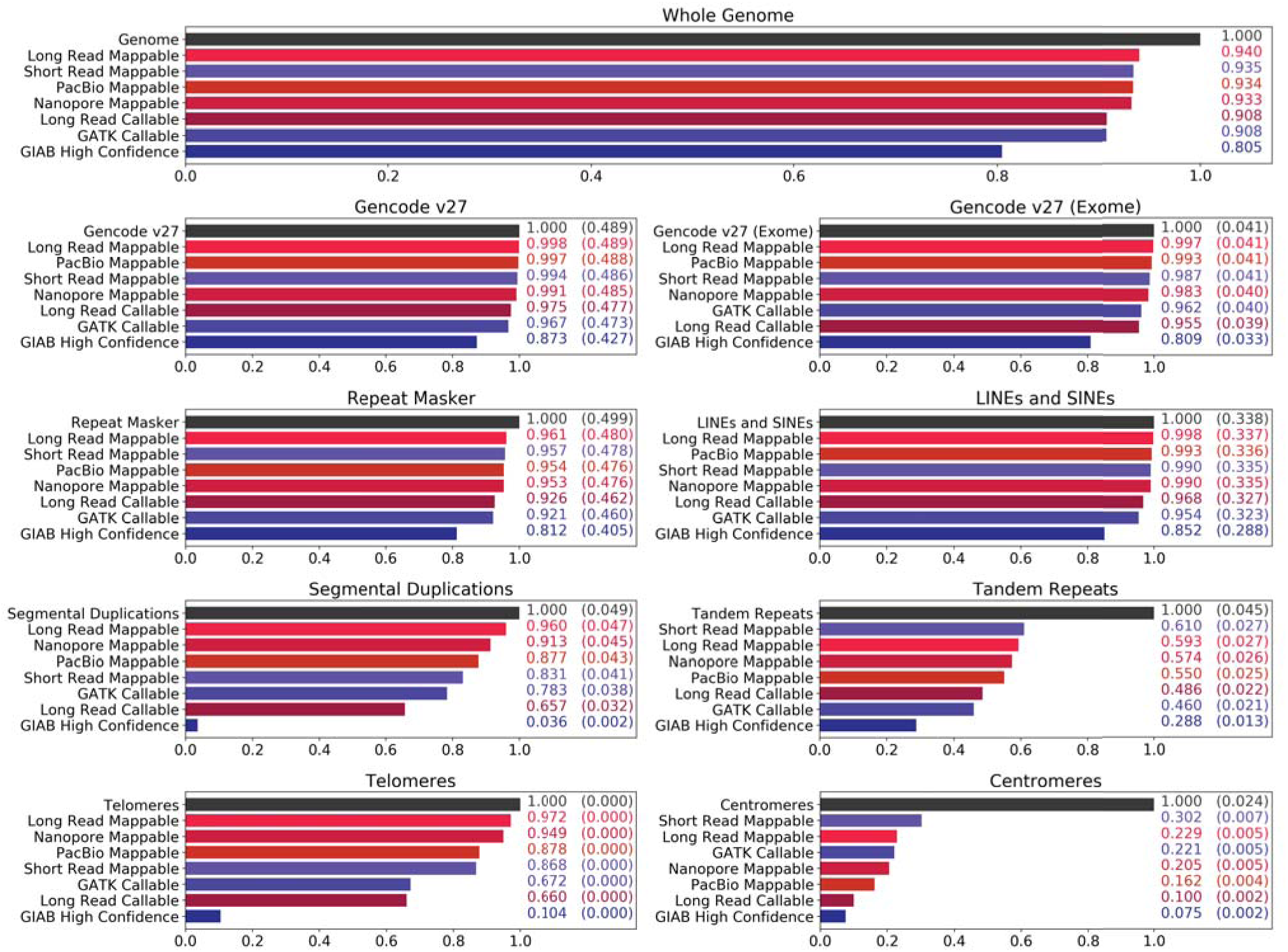
Reach of short read and long read technologies. The callable and mappable regions for NA12878 spanning various repetitive or duplicated sequences on GRCh38 is shown. Feature locations are determined based on BED tracks downloaded from the UCSC Genome Browser (21). Other than the Gencode regions (22, 23), all features are subsets of the Repeat Masker (24) track. Four coverage statistics for long reads (reds) and three for short reads (blues) are shown. PacBio and Nanopore describe areas where at least one primary read with GQ ≥ 30 has mapped, and Long Read Mappable describes where this is true for at least one of the long read technologies. Long Read Callable describes areas where both read technologies have coverage of at least 10 and less than twice the median coverage. GIAB High Confidence, GATK Callable and Short Read Mappable are the regions associated with the evaluation callsets. For the feature-specific plots, the numbers on the right detail coverage over the feature and (parenthesized) coverage over the genome.

Both sets of reads were aligned to GRCh38 with minimap2, a mapper designed to align error-prone long reads (27).

To ensure that any variants we found were not artifacts of misalignment, we filtered out reads flagged as secondary or supplementary, as well as reads with a mapping quality score less than 30. Genome-wide, this left approximately 12 million Nanopore reads and 34 million PacBio reads. The Nanopore reads had a median depth of 37× and length of 5950, including a set of ultra-long reads with lengths up to 900 kilobases. The PacBio reads had a median depth of 46× and length of 2650.

To validate the performance of our methods, we used callsets from Genome in a Bottle’s (GIAB) benchmark small variant calls v3.3.2 (10). First, we compared against GIAB’s set of high confidence calls, generated by a consensus algorithm spanning multiple sequencing technologies and variant calling programs. The high confidence regions associated with this callset exclude structural variants, centromeres, and heterochromatin. We used this to show our method’s accuracy in well-understood and easy-to-map regions of the genome.

We also analyzed our results compared to two callsets which were used in the construction of GIAB’s high confidence variants, one made by GATK HaplotypeCaller v3.5 (GATK/HC, 1) and the other by Freebayes 0.9.20 (28), both generated from a 300× PCR-free Illumina sequencing run (10).

All of our evaluation statistics were generated with the tool `vcfeval` from Real Time Genomics (29). We restrict the analysis to SNVs due to the error distribution of both PacBio and Nanopore long reads which leads to insertions and deletions being the most common type of sequencing error by far (30,31).

#### Short read variant callers

We explored the suitability of current state-of-the-art callers for short reads to process long read data (using default settings), but were unsuccessful. The absence of base qualities in the PacBio data prevented any calling; for Nanopore data, FreeBayes was prohibitively slow and neither Platypus nor GATK/HC produced calls.

### 3.2 Long Read Coverage

We determined the regions where long and short reads can be mapped to the human genome. In Figure 6, various coverage metrics for short and long reads are plotted against different genomic features, which were mostly selected for being repetitive or duplicated.

The callsets on the Illumina data made by GATK/HC and FreeBayes come with two BED files describing where calls were made with some confidence. The first, described in Figure 6 as *Short Read Mappable*, was generated using GATK CallableLoci v3.5 and includes regions where there is a) at least a read depth of 20, and b) at most a depth of twice the median depth, only including reads with mapping quality of at least 20. This definition of callable only considers read mappings. The second, described as *GATK Callable*, was generated from the GVCF output from GATK/HC by excluding areas with genotype quality less than 60. This is a more sophisticated definition of callable as it reflects the effects of homopolymers and tandem repeats. We use these two BED files in our analysis of how short and long reads map differently in various areas of the genome.

For long reads, we show four coverage statistics. The records marked as “Mappable” describe areas where there is at least one high quality long read mapping (PacBio, Nanopore, and *Long Read Mappable* for areas where at least one of the technologies mapped). The *Long Read Callable* entries cover a conservative region which has a sufficient read depth to illustrate the efficacy of our method; it covers regions where both sequencing technologies had a minimum depth of ten and maximum of 2 × the median depth (similar to the CallableLoci metric).

Figure 6 shows that in almost all cases, long reads map to more area than is mappable by short reads. For example, nearly half a percent of the genome is mappable by long reads but not short reads. Long reads also map to one percent more of the exome, and thirteen percent more of segmental duplications. Centromeres and Tandem Repeats are outliers to this generalization, where neither PacBio nor Nanopore cover appreciably more than Illumina.

### 3.3 Comparison Against High Confidence Truthset

To validate our method, we first analyzed the SNV detection and genotyping performance of our algorithm using the GIAB high confidence callset as a benchmark. All variants reported in these statistics fall within the GIAB high confidence regions.

Figure 7 (top) shows precision and recall of our algorithms on both the PacBio and Oxford Nanopore data sets. MarginPhase and WhatsHap perform similarly overall. MarginPhase achieved higher precision and recall on Nanopore reads, with precision of 0.7686 and recall of 0.8089, compared to WhatsHap’s precision of 0.7131 and recall of 0.7248 on the same set of Nanopore reads. WhatsHap obtained better results on PacBio data, with a precision of 0.9738 and recall of 0.9593, compared to MarginPhase’s precision of 0.9497 and recall of 0.9147.

**Fig. 7.**
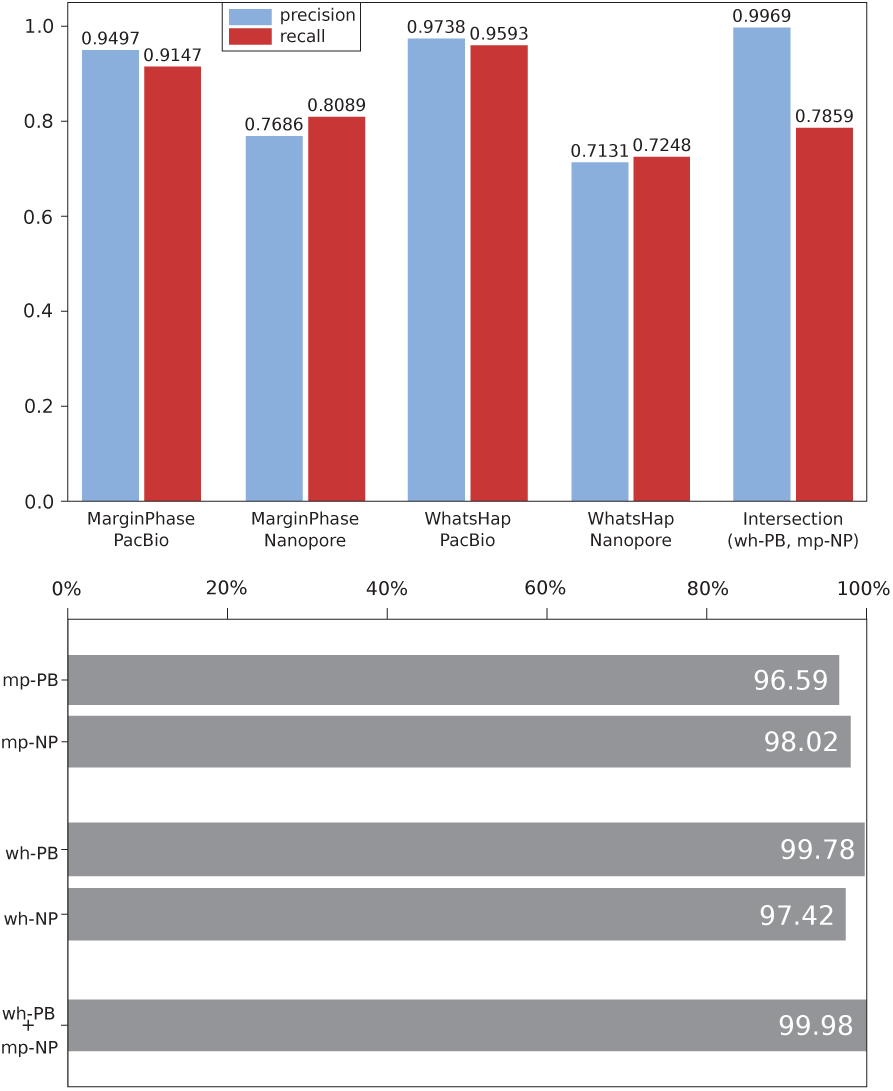
Precision and Recall (Top). of MarginPhase and WhatsHap on PacBio and Nanopore data sets in GIAB high confidence regions. **Genotype Concordance (Bottom)** (wrt. GIAB high confidence calls) of MarginPhase (mp, top) and WhatsHap (wh, middle) callsets on PacBio (PB) and Nanopore (NP) data. Furthermore, genotype concordance for the intersection of the calls made by WhatsHap on the PacBio and MarginPhase on the Nanopore reads is shown (bottom).

In addition to considering the two methods individually, we examine a combined set of variants which occur in both the calls made by WhatsHap on the PacBio reads and MarginPhase on the Nanopore data and where both tools report the same genotype. This improves the precision to 0.9969 at a recall of 0.7859. In further analysis, we refer to this combined variant set as *Long Read Variants*. It reflects a high precision subset of long read variants, validated independently by both sequencing technologies.

In order to further analyze the quality of the genotype predictions of our methods, we computed the genotype concordance of our callsets with respect to the GIAB ground truth inside of the high confidence regions. This was done by considering all variant positions correctly identified by MarginPhase and WhatsHap, and finding what fraction of these were also correctly genotyped (homozygous or heterozygous) with respect to the truth set. Figure 7 (bottom) shows the results. On the PacBio data, WhatsHap genotypes 99.78% of the variants contained in the truth set correctly, and MarginPhase genotypes 96.59% correctly. On the Nanopore data, MarginPhase performs slightly better by genotyping 98.02% of the SNVs contained in the GIAB callset correctly, while WhatsHap computed correct genotypes for 97.42% of the variants overlapping the GIAB truth set. Considering the intersection of the WhatsHap calls on PacBio, and MarginPhase calls on Nanopore data (i.e. our *Long Read Variants* set), we obtain a genotype concordance of 99.98%.

### 3.4 Cutting and Downsampling Reads

Our genotyping model incorporates haplotype information into the genotyping process by using the property that long sequencing reads can cover multiple variant positions. Therefore, one would expect the genotyping results to improve as the length of the provided sequencing reads increases. Furthermore, the coverage of the data would also affect the genotyping results.

In order to examine how the genotyping performance depends on the length of the sequencing reads and the coverage of the data, the following experiment was performed using the WhatsHap implementation. Both data sets (PacBio, Nanopore) were downsampled to average coverages 10 ×, 20 ×, 25 × and 30 ×. All SNVs inside of the high confidence regions in the GIAB truth set were re-genotyped from each of the resulting downsampled read sets, as well as from the full coverage data sets. Two versions of the genotyping algorithm were considered. First, the full length reads as given in the BAM files were provided to WhatsHap. Second, in an additional step prior to genotyping, the aligned sequencing reads were cut into shorter pieces such that each resulting fragment covered at most two variants. Additionally, we cut reads into fragments covering only one variant position. The genotyping performances of these genotyping procedures were finally compared by determining the amount of incorrectly genotyped variants.

Figure 8 shows the results of this experiment. On both data sets, the genotyping error increases as the length of reads decreases. Especially at lower coverages, the genotyping algorithm benefits from using the full length reads, which leads to much lower genotyping errors compared to using the shorter reads. In general, the experiment demonstrates that incorporating haplotype information gained from long reads does indeed improve the genotyping performance. Computing genotypes based on bipartitions of reads that represent possible haplotypes of the individual helps to reduce the number of genotyping errors, because it makes it easier to detect sequencing errors in the given reads.

**Fig. 8.**
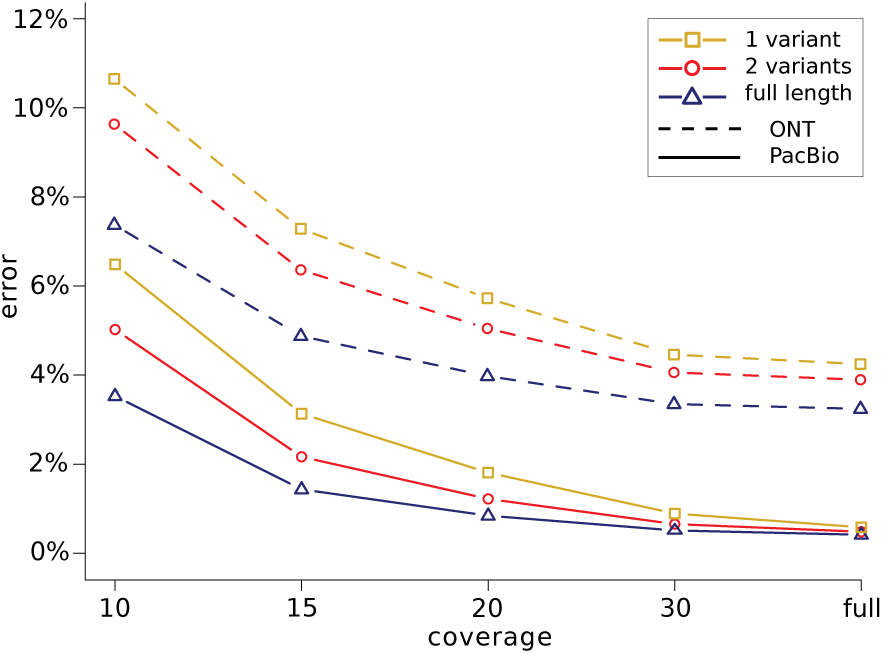
Genotyping Errors. (wrt. to GIAB calls) as a function of coverage. The full length reads were used for genotyping (blue) and additionally, reads were cut such as to cover at most two variants (red) and one variant (yellow). Solid lines correspond to PacBio, dashed lines to Nanopore data.

### 3.5 Callset Consensus Analysis

In Figure 9, we further dissect the relation of our intersection call set *(Long Read Variants,* which refers to variants called by both WhatsHap on PacBio reads and MarginPhase on nanopore reads) to the GIAB truth set, as well as to the callsets from GATK/HC and FreeBayes, which both contributed to the GIAB truth set.

**Fig. 9.**
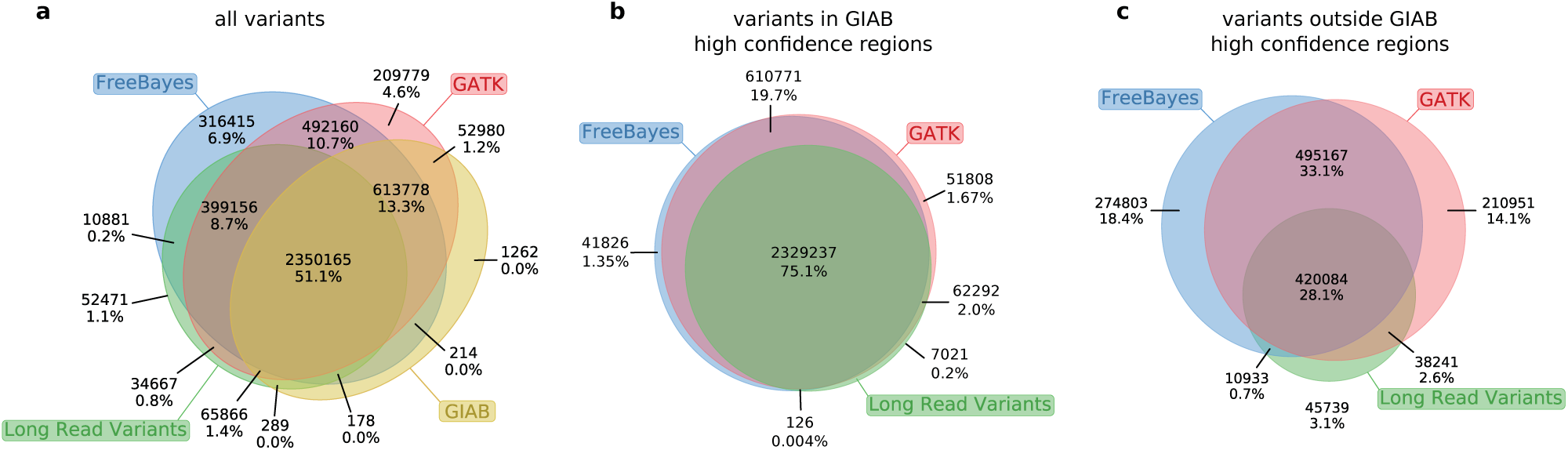
Confirming Short Read Variants. We examine all distinct variants found by our method, GIAB High Confidence, GATK/HC, and FreeBayes. Raw variant counts appear on top of each section, and the percentage of total variants is shown on bottom.

Figure 9a reveals that 399 156 variants in our *Long Read Variants* callset were called by both the GATK Haplotype Caller and FreeBayes, but are not in the GIAB truth set. To gather additional support for the quality of these calls, we consider two established quality metrics: the transition/transversion ratio (Ti/Tv), and the heterozygous/non-ref homozygous ratio (het/hom) (32). The Ti/Tv ratio of these variants is 2.10 and the het/hom ratio is 1.29. These ratios are comparable to those of the GIAB truth set, which are 2.10 and 1.55, respectively. An examination of the Platinum Genomes benchmark set (33), an alternative to GIAB, reveals 71371 such longread validated variants outside of their existing truth set.

We hypothesized that a callset based on long reads is particularly valuable in regions that were previously difficult to characterize. To investigate this, we separately examined the intersections of our *Long Read Variants* callset with the two short-read callsets both inside the GIAB high confidence regions and outside of them, see Figure 9b and Figure 9c, respectively. These Venn diagrams clearly indicate that the concordance of GATK and FreeBayes was indeed substantially higher in high confidence regions than outside. An elevated false positive rate of the short-read callers outside the high confidence regions is a plausible explanation for this observation. Interestingly, the fraction of calls concordant between FreeBayes and GATK for which we gather additional support is considerably lower outside the high confidence regions. This is again compatible with an increased number of false positives in the short read callsets, but we emphasize that these statistics should be interpreted with care in the absence of a reliable truth set for these regions.

### 3.6 Candidate Novel Variants

To demonstrate that our method allows for variant calling on more regions of the genome than short read variant calling pipelines, we have identified 15 498 variants which lie outside of the *Short Read Mappable* area, but inside the *Long Read Callable* regions, i.e. regions in which there is sequencing depth of at least 10 and not more than 2x the median depth for both sequencing technologies. We determined that 4.43 megabases of the genome (0.146%)is only mappable by long reads in this way.

Table 1 provides the counts of all variants found in each of the regions from Figure 6, as well as the counts for candidate variants, among the different types of genomic features described in Section 3.2. Over two thirds of the candidate variants occurred in the repetitive or duplicated regions described in the UCSC Genome Browser’s repeatMasker track. The transition/transversion ratio of NA12878’s 15 498 candidate variants is 1.64, and the heterozygous/homozygous ratio of these variants is 0.31. Given that we observe one candidate variant in every 325 haplotype bases, compared to one variant in every 1151 haplotype bases in the GIAB truth set, these candidate variants exhibit a 3.6 × increase in the haplotype variation rate.

**Table 1.** Distribution of candidate novel variants across different regions of interest. All variants refers to the variants in the Long Read Variants set, and Novel Variant Candidates are those described in Section 3.6.

|  | All Variants | Novel Variant Candidates |
| --- | --- | --- |
| **Total** | 2,913,942 | 15,498 |
| Genecode v27 (ALL) | 1,363,064 | 5,594 |
| Genecode v27 exome | 86,357 | 538 |
| Repeat Masker | 1,583,684 | 10,677 |
| LINEs | 690,859 | 5,161 |
| SINEs | 421,340 | 1,432 |
| Segmental Duplications | 157,341 | 5,683 |
| Tandem Repeats | 96,871 | 5,437 |
| Centromeres | 18,644 | 2,031 |
| Telomeres | 295 | 14 |

### 3.7 Runtimes

Whole genome variant detection using WhatsHap took 147 CPU hours on PacBio reads and 79.5 hours on Nanopore, of which genotyping took 42.2 and 32.8 hours respectively. The MarginPhase implementation took 583 CPU hours on PacBio and 330 on Nanopore, with an additional 1730 and 1220 hours for realignment.

## 4 Discussion

We present a method that uses a Hidden Markov Model to partition long reads into haplotypes, which we found to improve the quality of variant calling. This is evidenced by our experiment in cutting and downsampling reads, where reducing the number of variants spanned by any given read leads to decreased performance at all levels of read coverage.

Our analysis of the method against a high confidence truth set in high confidence regions shows false discovery rates (corresponding to one minus precision) between 3 and 6 percent for PacBio, and between 24 and 29 percent for Nanopore. However, when considering a conservative set of variants confirmed by both long read technologies, the false discovery rate drops to around 0.3%, comparable with contemporary short read methods in these regions.

In analyzing the area of the genome with high quality long read mappings, we found roughly a half a percent of the genome (approximately fifteen megabases) that is mappable by long reads but not by short reads. This includes one percent of the human exome, as well as over ten percent of segmental duplications. Even though some of these areas have low read counts in our experimental data, the fact that they have high quality mappings means that they should be accessible with sufficient sequencing. We note that this is not the case for centromeric regions, where Illumina reads were able to map over twice as much as we found in our PacBio data. This may be a result of the low quality in long reads preventing them from uniquely mapping to these areas with an appreciable level of certainty.

Over our entire set of called variants, the Ti/Tv and het/hom ratios were similar to those reported by the truth set. The Ti/Tv ratio of 2.18 is slightly above the 2.10 reported in the GIAB callset, and the Het/Hom ratio of 1.36 is lower than the 1.55 found in the GIAB variants. In the 15 498 novel variant candidates produced by our method in regions unmappable by short reads, the Ti/Tv ratio of 1.64 is slightly lower than that of the truth set. This is not unexpected as gene-poor regions such as these tend to have more transversions away from C:G pairs (34). We also observe that the Het/Hom ratio dropped to 0.31, which could be due to systematic biases in our callset or in the reference genome. The rate of variation in these regions was also notably different than in the high confidence regions, where we find three variants per thousand haplotype bases (3.6× the rate in high confidence regions). A previous study analyzing NA12878 (35) also found an elevated variation rate in regions where it is challenging to call variants, such as low complexity regions and segmental duplications. The study furthermore found clusters of variants in these regions, which we also observe.

The high precision of our intersected Nanopore/PacBio long read variants set makes it useful as strong evidence for confirming existing variant calls. As shown in the read coverage analysis, in both the GIAB and Platinum Genomes efforts many regions cannot be called with high confidence. In the excluded regions of GIAB we found just under 400 thousand variants using both Nanopore and PacBio reads with our methods, which were additionally confirmed with Illumina reads by two other variant callers, FreeBayes and GATK/HC. Given the extensive support of these variants from multiple sequencing technologies and variant callers, these variants are good candidates for addition to the GIAB truth set. Expansion of benchmark sets to harder-to-genotype regions of the human genome is generally important for the development of more comprehensive genotyping methods, and we plan to work with these efforts to use our results. Further, our method is likely to prove useful for future combined diplotyping algorithms when both genotype and phasing is required, for example as may be used when constructing phased diploid *de novo* assemblies (36) or in future hybrid long/short read diplotyping approaches.

## Acknowledgements

We thank the GIAB project for providing the data sets used. In particular, we thank Justin Zook for helpful discussions on how to use GIAB data, and Miten Jain for help with the nanopore data. This work was supported, in part, by the National Human Genome Research Institute of the National Institutes of Health under Award Number 5U54HG007990 and grants from the W.M. Keck foundation and the Simons Foundation.

